# Novel genetically encoded tools for imaging or silencing neuropeptide release from presynaptic terminals *in vivo*

**DOI:** 10.1101/2023.01.19.524797

**Authors:** Dong-Il Kim, Sekun Park, Mao Ye, Jane Y. Chen, Jinho Jhang, Avery C. Hunker, Larry S. Zweifel, Richard D. Palmiter, Sung Han

## Abstract

Neurons produce and release neuropeptides to communicate with one another. Despite their profound impact on critical brain functions, circuit-based mechanisms of peptidergic transmission are poorly understood, primarily due to the lack of tools for monitoring and manipulating neuropeptide release *in vivo*. Here, we report the development of two genetically encoded tools for investigating peptidergic transmission in behaving mice: a genetically encoded large dense core vesicle (LDCV) sensor that detects the neuropeptides release presynaptically, and a genetically encoded silencer that specifically degrades neuropeptides inside the LDCV. Monitoring and silencing peptidergic and glutamatergic transmissions from presynaptic terminals using our newly developed tools and existing genetic tools, respectively, reveal that neuropeptides, not glutamate, are the primary transmitter in encoding unconditioned stimulus during Pavlovian threat learning. These results show that our sensor and silencer for peptidergic transmission are reliable tools to investigate neuropeptidergic systems in awake behaving animals.

## INTRODUCTION

Two types of transmitters are found within synaptic terminals for neuronal communication: classical neurotransmitters and neuromodulators. Classical fast-acting neurotransmitters, including glutamate, acetylcholine, GABA, and glycine, are packaged into synaptic vesicles (SVs), which are released from synaptic terminals in an activity-dependent manner (Sudhof, 2012). By contrast, neuromodulators, such as neuropeptides and monoamines, are stored in large dense-core vesicles (LDCVs), which are released in response to a train of high-frequency action potentials (Cifuentes et al., 2008; Martinez-Rodriguez and Martinez-Murillo, 1994; Salio et al., 2006; Silm et al., 2019). The unique functions and distinct release properties of neuromodulators suggest that they can function as the main neurotransmitter in neurons, like dopamine and norepinephrine (Eskenazi et al., 2021; Hnasko and Edwards, 2012; Vaaga et al., 2014). Neuropeptides are by far the most diverse class of neuromodulators. Currently, more than 100 neuropeptides and their postsynaptic receptors have been discovered, with each neuropeptide exhibiting unique functions, such as arousal, sleep/wake, reproduction, feeding, reward, learning/memory, and threat perception (van den Pol, 2012). Furthermore, dysregulation of neuropeptides has been closely associated with many neurological and neuropsychological disorders (Beal and Martin, 1986). Thus, elucidating the mechanism by which neuropeptidergic systems act in brain circuits is critical for understanding brain function and associated disorders, and it requires the tools that monitor and manipulate peptidergic transmissions in a temporally precise manner in behaving animals. To date, this is a technical feat with very limited availability.

Recent progress in developing neuromodulator sensors by genetically modifying their postsynaptic G-protein-coupled receptors (GPCR) allow researchers to monitor the release of many monoamine and catecholamine neuromodulators in behaving animals (Sabatini and Tian, 2020; Wu et al., 2022). The development of neuropeptide sensors is also progressing rapidly (Melzer et al., 2021; Qian et al., 2022; Wu et al., 2022). However, it is impossible to develop a universal neuropeptide sensor by engineering postsynaptic GPCRs due to their diversity. Furthermore, despite their fundamental contribution on systemic understanding of peptidergic circuits in the brain, the postsynaptic GPCR-based neuropeptide sensors has some inherent limitations primarily because their site of action is postsynaptic (Rusakov, 2022). To resolve these issues, several attempts have been made to develop presynaptic neuropeptide sensors by genetically labeling neuropeptides with a fluorescent protein (Ding et al., 2019; Shaib et al., 2018; Taraska et al., 2003). However, this approach has never been applied to awake behaving animals, because fluorescently labeled neuropeptides are rapidly depleted after release. Here, we report a new genetically encoded fluorescent LDCV release sensor that directly monitors the release of neuropeptides from presynaptic terminals in behaving mice. The sensor uses a pH-sensitive variant of GFP, superecliptic pHluorin (SEP) (Sankaranarayanan et al., 2000), to detect pH changes inside the lumen of LDCVs during release events. We targeted SEP to the luminal membrane of the LDCV by incorporating it into the luminal loop of cytochrome b561 (CYB561), a LDCV-specific membrane protein (Birinci et al., 2020; Perin et al., 1988). We validated and optimized the CYB561-SEP fusion protein as a LDCV sensor (CybSEP) in differentiated PC12 pheochromocytoma cell lines, mouse brain slices, and awake behaving mice.

Peptidergic neurons co-express fast-acting transmitters and neuropeptides, and they are packaged in different vesicles that have distinct release properties suggesting that each transmitter may shape the function of these neurons differentially. However, circuit-specific dissection of functional roles played by each transmitter is impossible due to the lack of tools that specifically manipulate peptidergic transmissions leaving fast neurotransmission unaltered. Here, we report a novel genetically encoded silencer that specifically blocks peptidergic transmissions without changing fast neurotransmission. The silencer uses a neuropeptide-specific peptidase, neutral endopeptidase (NEP), also called neprilysin or enkephalinase, which specifically inactivates many neuropeptides, including but not limited to enkephalin, bradykinin, calcitonin gene-related peptide (CGRP), substance P, neurotensin, and oxytocin by cleaving their hydrophobic amino acid chains (Gourlet et al., 1997; Katayama et al., 1991; Scholzen and Luger, 2004; Skidgel et al., 1984; Stancampiano et al., 1991; Turner et al., 1985a, b). We targeted the NEP to the luminal LDCV by combining it with the LDCV-targeting signal peptide, then validated this LDCV-targeted NEP as a silencer of peptidergic transmission electrophysiologically in mouse brain slices, and behaviorally in awake behaving mice. Our study, using novel neuropeptide release sensor and silencer, demonstrates that neuropeptides, not glutamate, are shown to be essential for conveying aversive unconditioned stimuli during Pavlovian threat learning.

## RESULTS

### Design and characterization of the LDCV sensor

SEP was originally engineered to detect synaptic transmission by targeting it to the inside of SVs by incorporating it into SV-specific membrane proteins, such as synaptobrevin or synaptophysin (Miesenböck et al., 1998; Zhu et al., 2009). We re-purposed this proven system to monitor LDCV release by fusing SEP to CYB561. The CYB561 is a transmembrane electron transport protein unique to LDCVs (Fleming and Kent, 1991; Perin et al., 1988). It mediates transmembrane electron transport to regenerate ascorbic acid inside the lumen of LDCVs and plays a critical role in the biosynthesis of neuropeptides by supplying reducing equivalents to α-amidase (Lu et al., 2014). We first substituted two histidines for alanines at positions 86 and 159. These histidines are essential sites for ascorbate binding, and therefore replacing them with alanines eliminated CYB561’s electron transporter activity (Kipp et al., 2001; Lu et al., 2014). We then inserted SEP coding sequence into the luminal domain of CYB561, between transmembrane domain 3 and 4 (CybSEP) (Figure 1A). In addition, we created a control fusion protein by inserting gamillus, an acid-tolerant monomeric GFP, into the same position of Cyb561 (CybGam) (Shinoda et al., 2018). To determine whether CybSEP alters fluorescence in response to changes in pH, we expressed CybSEP in PC12 cells differentiated by nerve growth factor and perfused acidic solution (pH 5.5) followed by NH_4_Cl treatment to deacidify intracellular compartments. We observed almost a complete loss of CybSEP fluorescence in the acidic condition. The signal returned following NH_4_Cl perfusion. In contrast, CybGam showed no changes in fluorescence (Figures 1B-1D) under the same conditions. Bath application of 70 mM KCl, which leads to membrane depolarization, induced robust increases in CybSEP fluorescence, whereas CybGam exhibited no detectable changes in fluorescence in response to KCl (Figures 1E, 1F). We further engineered the CybSEP to increase fluorescence intensity by incorporating two SEPs into CYB561 (CybSEP2). Evoked changes in fluorescence were 30% larger for CybSEP2 compared with CybSEP (Figures S1A and S1B). Thus, CybSEP2 was used for the rest of the experiments.

**Figure 1.**
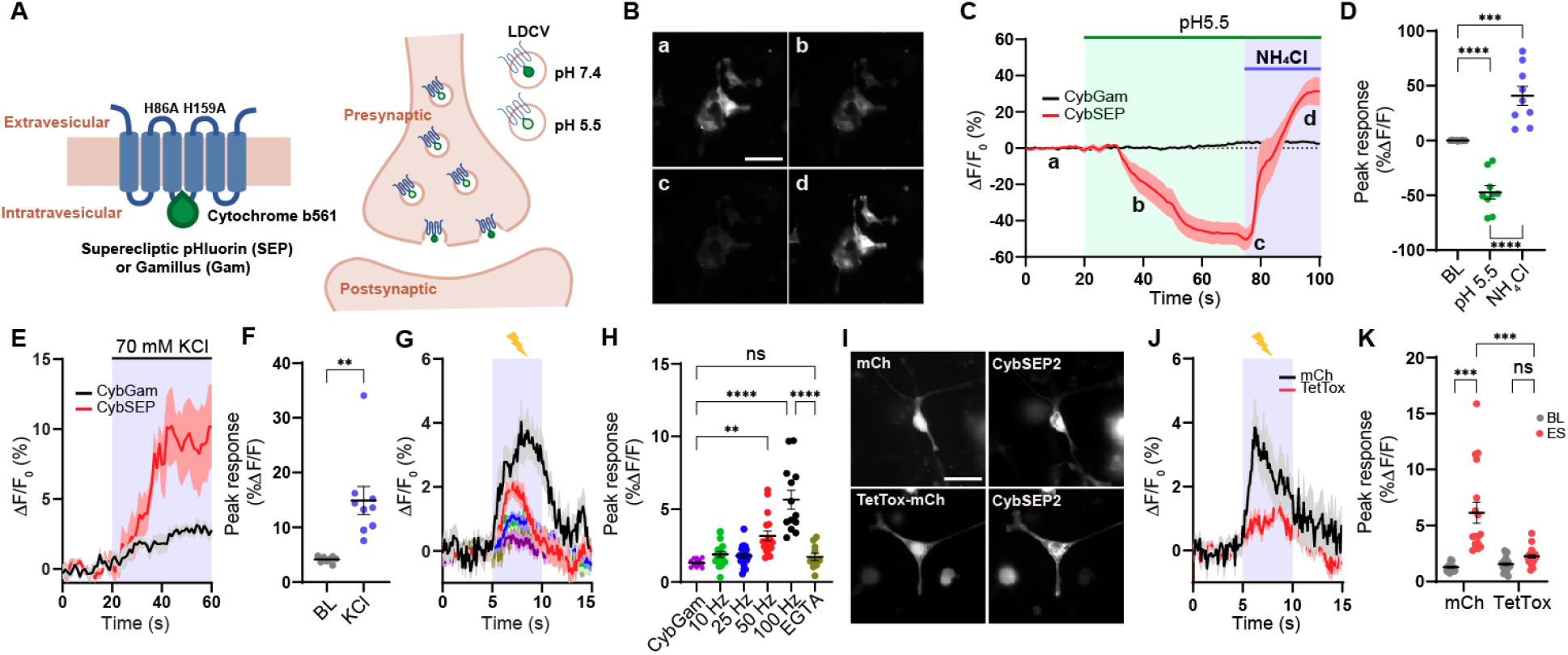
Design and characterization of CybSEP as an LDCV sensor. (A) Schematic of constructs and working principle. (B) Representative images of CybSEP expression in differentiated PC12 cells when perfused with acidic and NH_4_Cl solutions, described in (C) (a, bath solution; b and c, acidic solution; d, NH_4_Cl; Scale bar, 100 µm). (C) The traces of CybSEP and CybGam fluorescence change during application of various extracellular solutions. (D) Quantification of percent ΔF/F_0_ peak intensity in CybSEP expressing PC12 cells (n= 9 over 3 experimental replicates; ***p < 0.001, ****p < 0.0001 via one-way ANOVA followed by Tukey’s multiple comparisons). BL indicates baseline. (E and F) Average trace of fluorescence change during 70 mM KCl treatment and quantification of percent ΔF/F_0_ peak intensity in (E) (n=8-10 over 2 experimental replicates; **p < 0.01 via paired t test to the baseline). (G and H) Average traces of fluorescence change during various electrical stimulation and quantification of percent ΔF/F_0_ peak intensity in (G). For extracellular calcium removal, 5 mM EGTA was used instead of CaCl_2_ (n=11-19, over 3 experimental replicates; **p < 0.001, ****p < 0.0001 via one-way ANOVA followed by Tukey’s multiple comparisons). (I) Representative images of CybSEP2 with mCherry (mCh) or TetTox-mCh expressed in PC12 cells (Scale bar, 100 µm). (J and K) Average traces of fluorescence change in CybSEP2 co-expressed with mCh or TetTox-mCh during electrical stimulation at 100 Hz and quantification of percent ΔF/F_0_ peak intensity in (J) (n=19-23, 3 over 3 experimental replicates; ***p < 0.0001 via two-way ANOVA followed by Šidák multiple comparisons). Data are represented as mean ± SEM.

Electrical stimulation of differentiated PC12 cells at various frequencies (10, 25, 50, and 100 Hz) via a glass pipet showed frequency-depended increases in fluorescent signal (the maximum response was observed at 100 Hz). Moreover, EGTA treatment abolished the fluorescent signal evoked by 100-Hz stimulation, indicating that the response depended on calcium (Figures 1G-1H) (Nakamura, 2019). To test whether the fluorescent response resulted from LDCV release events, we inhibited vesicular fusion using the tetanus toxin light chain (TetTox), which disrupts the release of both SVs and LDCVs by cleaving synaptobrevin (McMahon et al., 1992). Expressing TetTox-mCherry with CybSEP2 resulted in a greatly reduced fluorescent signal in response to electrical stimulation (100 Hz), compared with mCherry expressing controls (Figures 1I-1K). These data indicate that fluorescent responses from CybSEP2 reflect LDCV release events.

### Imaging presynaptic LDCV release in acute brain slices

After successfully validating that CybSEP2 can be used to detect LDCV release in the differentiated PC12 cells, we tested it in intact brain slices. The parabrachio-amygdaloid pathway is a well-described peptidergic circuit that encodes aversive unconditioned sensory stimuli (US) during Pavlovian threat learning (Nagase et al., 2019). Previous studies have shown that peptidergic neurons expressing CGRP (encoded by the *Calca* gene) in the external lateral parabrachial nucleus (PBel) and their direct downstream CGRP receptor-expressing neurons in the lateral subdivision of the central amygdala (CeAl) plays important roles in pain perception and threat learning in mice (Han et al., 2005; Han et al., 2015; Salmon et al., 2001; Sato et al., 2015). In addition to glutamate, CGRP neurons in the PBel (CGRP^PBel^) co-express various neuropeptides, such as substance P (SP), pituitary adenylyl cyclase-activating peptide (PACAP), and neurotensin (NTS) (Kang et al., 2020; Palmiter, 2018; Pauli et al., 2022). However, the main transmitter that relays the US information in this peptidergic circuit was not known. Therefore, this circuit is ideal for functionally validating CybSEP2 in the brain. To monitor peptidergic transmissions in this circuit, recombinant adeno-associated viruses (AAVs) Cre-dependently encoding CybSEP2 or CybGam (AAV_DJ_-DIO-CybSEP2 or AAV_DJ_-DIO-CybGam) were bilaterally injected into the PBel of *<u>Calca</u>*^Cre/+^ mice (Figures 2A and S2A). Four weeks after the injection, we confirmed that CybSEP2 and CybGam were robustly expressed in neuronal cell bodies within the PBel, and in axons projecting to the CeAl (Figures 2B and S2A). A high-speed fluorescence imaging system was used to monitor the CybSEP2 fluorescence changes in acutely dissociated brain slices. Electrical stimulation of CGRP^PBel^ axonal terminals in the CeAl via a glass pipet resulted in frequency-dependent increases in fluorescence, with a maximum response at 100 Hz. These responses were abolished by bath application of calcium chelator, EGTA. No significant increases in fluorescence were observed in brain slices expressing CybGam (Figures 2C-2E). The rise time (τ_on_) of these responses was faster than the decay time (τ_off_) at 50 and 100 Hz (Figure 2F). Interestingly, significant fluorescence was also detected near the CGRP^PBel^ neuronal cell bodies and dendrites that express CybSEP2 (Figures S2B-S2D), an intriguing observation that the CGRP signaling might be functionally relevant in the PBel (Ludwig and Leng, 2006; van den Pol, 2012). Repeated stimulation of CGRP^PBel^ axonal terminals in the CeAl 4 times over 20 min with 5-min intervals evoked similar fluorescence signal, indicating that the CybSEP2 fluorescence signal was not depleted nor quenched under these conditions (Figures 2G and 2H). Inhibiting vesicular release by co-expressing TetTox in CGRP^PBel^ neurons of the *<u>Calca</u>*^Cre/+^ mice substantially attenuated 100 Hz-evoked changes in fluorescence in their axonal terminals within the CeAl (Figures 2I and 2J). By contrast, no noticeable decreases in fluorescence were observed in mCherry-expressing control brain slices (Figures 2K and 2L). Taken together, these results suggest that CybSEP2 can be used to reliably detect LDCV release events at axonal terminals in acutely dissociated brain slices, making it a useful tool to study neuropeptide release from presynaptic terminals.

**Figure 2.**
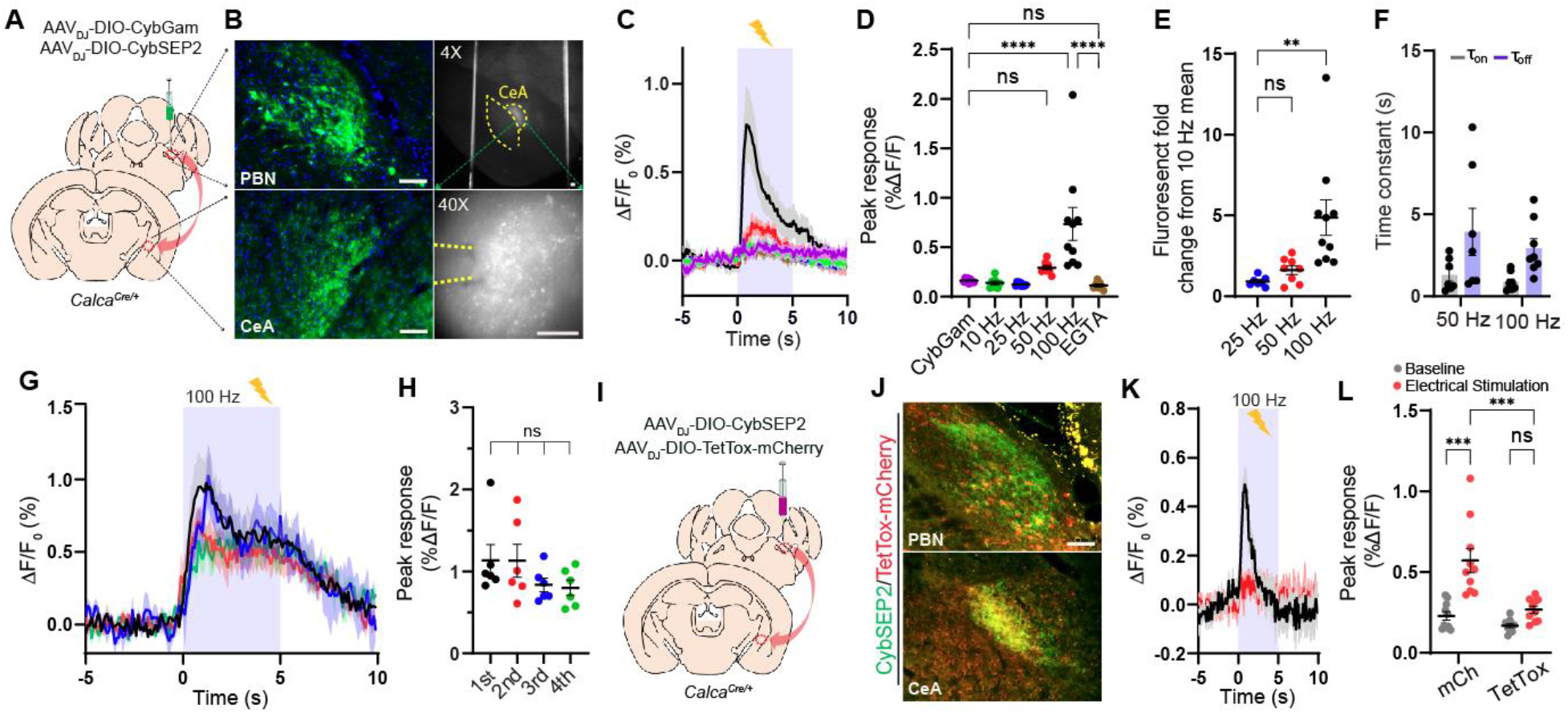
Imaging the LDCV release in brain slices. (A and B) Schematic and representative images showing CybSEP2 targeted region, its projection to the CeA and expression of CybSEP2 in the PBN and the CeA of the *Calca*^*Cre*/+^ brain slices. Images in the right panel (B) show slices containing the CeA for slice imaging experiment (Scale bar, 100 µm). (C and D) Average traces of fluorescence change in response to various electrical stimulation and quantification of the data (C). For extracellular calcium removal, 5 mM EGTA was used instead of CaCl_2_. Each trace is the average of 7-9 trials in 24 slice slices prepared from 4 mice (****p < 0.0001 via one-way ANOVA followed by Šidák multiple comparisons). (E) Quantification of fold change from percent ΔF/F_0_ peak intensity compared to 10 Hz in (D) (**p < 0.01 via one-way ANOVA followed by Tukey’s multiple comparisons). (F) Time constant (τ) of CybSEP2 expressing neurons during electrical stimulation at 50 Hz (n=7) and 100 Hz (n=8). The rising (τ_on_) and decay (τ_off_) phases were determined by fitting across an entire stimulation period (τ_on_ = 1.30 ± 0.37, τ_off_ = 3.93 ± 1.44 at 50 Hz; τ_on_ = 0.85 ± 0.18, τ_off_ = 2.93 ± 0.55 at 100 Hz). (G and H) Average traces of fluorescence change in response to repeated electrical stimulation at 100 Hz and quantification of percent ΔF/F_0_ peak intensity in (G). Each trial was measured at 5 min interval between trials (n=6; ns, not significant via one-way ANOVA followed by Tukey’s multiple comparisons). (I and J) Schematic brain region targeted for viral injection and co-expression of CybSEP2 and TetTox-mCherry in the PBN and in the CeA of *Calca*^*Cre/+*^ (Scale bar, 100 µm). (K and L) The trace of fluorescence change in CybSEP2 with mCherry (n=10 slices from 3 mice) or TetTox-mCherry (n=10 slices from 3 mice) expressing neurons during electrical stimulation at 100 Hz and quantification of date in (K) (***p < 0.0001 via two-way ANOVA followed by Šidák multiple comparisons). Data are represented as mean ± SEM.

### Monitoring presynaptic LDCV release in freely moving mice

The CGRP^PBel^ neurons are activated by unconditioned stimulus (electric foot shock), but not by the conditioned stimulus (tone) during Pavlovian threat conditioning (Kang et al., 2022). We therefore examined whether CybSEP2 can be used as a LDCV sensor to detect electric foot shock-evoked neuropeptide release from CGRP^PBel→CeAl^ terminals by monitoring fluorescence changes in the CeAl of freely moving mice during threat conditioning. We stereotaxically injected AAV_DJ_-DIO-CybSEP2 or AAV_DJ_-DIO-CybGam into the PBel of *Calca*^Cre/+^ mice, and then implanted a fiberoptic cannula into the CeAl (Figure 3B). A CMOS fiber photometry system was used to monitor LDCV release in freely moving mice (Figure 3A). Following 4 weeks of recovery from the stereotaxic surgery, CMOS fiber photometry revealed that a mild electric footshock (0.3 mA, 2 s) triggered a sharp increase in fluorescence in the CGRP^PBel→CeAl^ terminals of CybSEP2-expressing *Calca*^Cre/+^ mice, but not in CybGam-expressing control mice (Figures 3C and 3D). Since the CGRP^PBel^ neurons are also activated by painful stimuli, we monitored the neuropeptide releases evoked by noxious heat. In the hot plate test, 52°C thermal stimulus significantly increased the fluorescence signal in CGRP^PBel→CeAl^ terminals, whereas 42°C thermal stimulus did not. No changes in fluorescence were observed in response to both thermal stimuli (42 and 52°C) in CybGam-expressing mice (Figures 3E and 3F). We also tested quinine as another aversive sensory stimulus, since previous studies have shown that the CGRP^PBel→CeAl^ pathway is activated by quinine consumption (Kang et al., 2022). Fluorescence intensity rapidly increased at the onset of quinine consumption in CGRP^PBel→CeAl^ terminals of CybSEP2-expressing *Calca*^Cre/+^ mice, but not in CybGam-expressing control mice. Water consumption failed to evoke fluorescence changes in both groups of mice (Figures 3G and 3H). Taken together, CybSEP2 imaging in freely moving mice shows that aversive sensory stimuli induce robust signals from the LDCV sensor, indicative of neuropeptide release.

**Figure 3.**
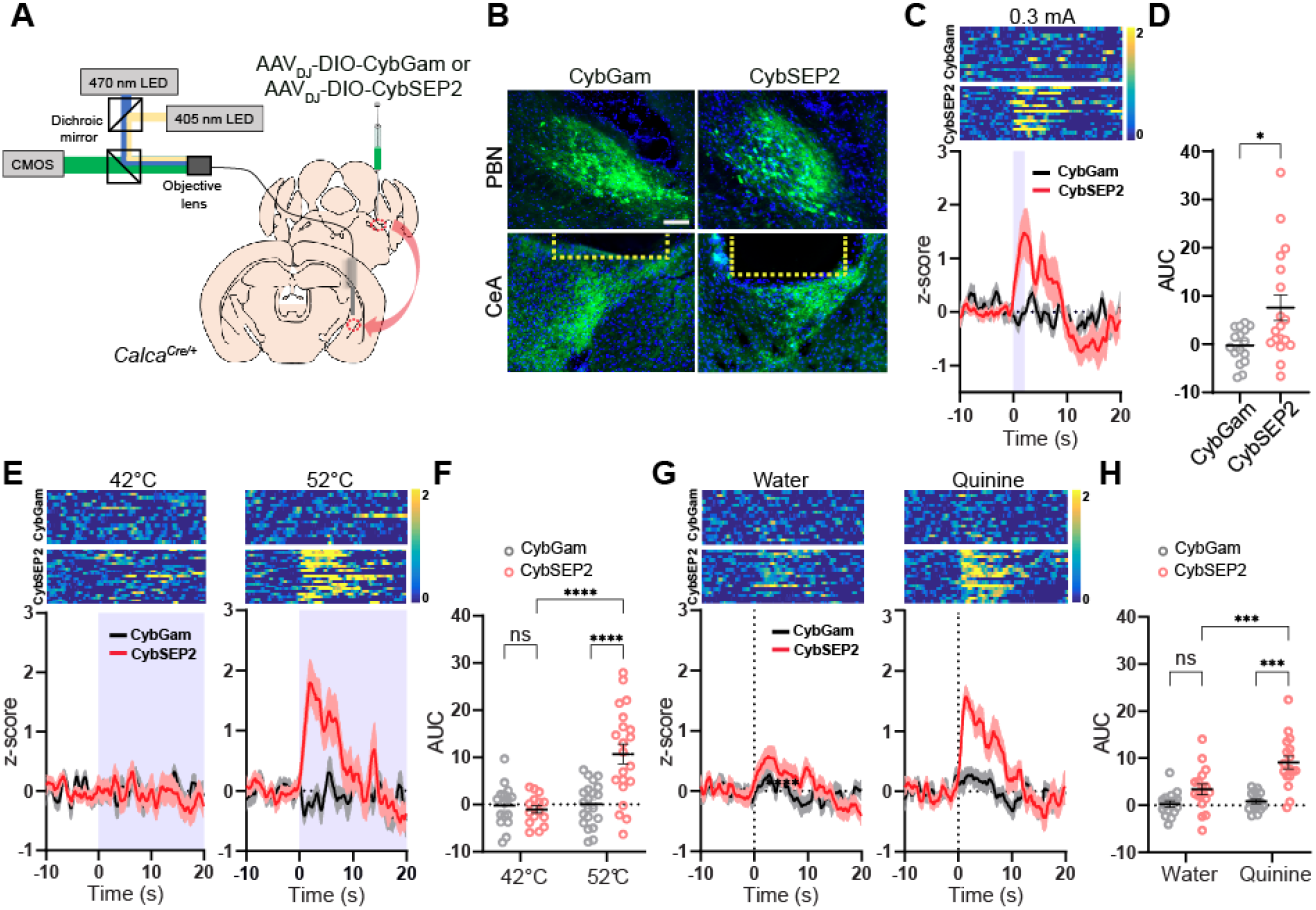
Monitoring LDCV release from the synaptic terminals in behaving mice. (A) Schematic illustration of fiber photometry system used for CybSEP2 response recording in the *Calca*^*Cre/+*^ mice. (B) Expression images of CybGam and CybSEP2 in the PBN and the CeA of *Calca*^*Cre/+*^ mice with an optic fiber implanted over the CeA. Yellow dot line represents the location of optic fiber (Scale bar, 100 µm). (C) Heat map and average traces of fluorescence change elicited by footshock (0.3 mA) in CybGam (16 traces from 4 mice) and CybSEP2 (18 traces from 5 mice) expressing mice. (D) Quantification of data in (C) by area under curve (AUC) for 0-10 s (*p < 0.005 via unpaired t-test comparisons to control). (E) Heat map and average traces of fluorescence change during thermal stimulus in CybGam (16 traces from 4 mice) and CybSEP2 (21 traces from 5 mice) expressing mice at 42 or 52°C hot plate with a cutoff time of 20 s. (F) Quantification of data in (E) by AUC for 0-10 s (****p < 0.0001 via two-way ANOVA followed by Tukey’s multiple comparisons). (G) Heat map and average traces of fluorescence change during aversive taste stimulus in CybGam (16 traces from 4 mice) and CybSEP2 (17 traces from 5 mice) expressing mice elicited by 0.5 mM quinine solution or water. (H) Quantification of data in (G) by AUC for 0-10 s (***p < 0.001via two-way ANOVA followed by Tukey’s multiple comparisons to the CybGam or CybSEP2). Data are represented as mean ± SEM.

CGRP^PBel^ neurons express multiple neuropeptides, but they also express vesicular glutamate transporter type 2 (Vglut2) and make glutamatergic synapses onto neuronal targets in the CeAl (Carter et al., 2013; Huang et al., 2021). To test how glutamatergic and peptidergic transmissions interact in CGRP^PBel→CeAl^ terminals in response to aversive sensory stimuli, we monitored fast neurotransmitter release from axonal terminals in response to the aversive sensory stimuli. To monitor glutamatergic transmissions in CGRP^PBel→CeAl^ terminals, we fused the SEP with the SV-specific protein, synaptophysin (SypSEP), and packaged it into an AAV vector with a DIO cassette (AAV_DJ_-DIO-SypSEP) (Granseth et al., 2006; Zhu et al., 2009). We expressed SypSEP in the PBel of *Calca*^Cre/+^ mice (Figure S3A) and examined fluorescence response evoked by various aversive stimuli. In contrast to CybSEP, SypSEP fluorescence did not increase, but rather slightly decreased from the baseline when mice were exposed to aversive sensory stimuli (Figures S3B-S3D). However, the SypSEP fluorescence was robustly increased as these mice consumed high-nutrient liquid formula (Ensure, Abbott) (Figure S3E). These results indicate that SEP-based LDCV or SV release sensors can detect peptidergic and glutamatergic transmissions independently, and together they can be used to distinguish between behaviors that result from LDCV vs. SV transmissions from presynaptic terminals of peptidergic neurons in freely moving mice.

### Inhibiting peptidergic transmission attenuates threat learning

Monitoring LDCV and SV releases from the CGRP^PBel→CeAl^ terminals showed that aversive sensory stimuli triggers the release of neuropeptides. To investigate whether neuropeptides play a pivotal role in transmitting aversive sensory stimuli to the amygdala during threat learning, we engineered a peptidase that selectively degrades neuropeptides by targeting it specifically into the luminal side of the LDCV. Neutral endopeptidase (NEP) is a transmembrane protease present at the cell surface which cleaves a broad range of neuropeptides that contain hydrophobic amino acid in the extracellular space (Helin et al., 1994; Hui, 2007; Katayama et al., 1991). The CGRP^PBel^ neurons co-express multiple neuropeptides including CGRP, neurotensin, substance P, and pituitary adenylate cyclase-activating peptide (Kang et al., 2020), all of which can be degraded by the NEP (Gourlet et al., 1997; Katayama et al., 1991; Skidgel et al., 1984). Therefore, we utilized the NEP to degrade active neuropeptides packaged inside the LDCVs of the CGRP^PBel^ neurons. We incorporated the LDCV targeting signal peptide from pro-opiomelanocortin (POMC) into the NEP to selectively translocate it to the luminal side of LDCV (Cool et al., 1995). We then constructed P2A-mediated bicistronic AAV vector encoding LDCV-targeted NEP (NEP_LDCV_), as well as the cytosolic mRuby3 (AAV_DJ_-DIO-NEP_LDCV_-P2A-mRuby3) (Figure 4A). To validate its expression in the mouse brain, AAV_DJ_-DIO-NEP_LDCV_-P2A-mRuby3 or AAV_DJ_-DIO-mCherry was bilaterally delivered into the PBel of *<u>Calca</u>*^*Cre/+*^ mice (Figure 4B). After confirming the expression of mRuby3 in the CGRP^PBel^ neurons, we immunostained the PBel-containing coronal brain sections with antisera against CGRP to evaluate the degradation of CGRP in the presence of the NEP_LDCV_. We found that most mCherry-expressing neurons were co-labeled with CGRP-immunofluorescent signals while CGRP-immunoreactivity were barely detected in NEP_LDCV_-expressing neurons suggesting efficient proteolytic degradation of CGRP by the NEP_LDCV_ (Figure 4C and 4D). We next sought to determine whether the loss of neuropeptides in the CGRP^PBel^ neurons affects peptidergic transmission in CGRP^PBel→CeAl^ synapses by electrophysiological recording of the postsynaptic CeAl neurons in brain slices. AAVs Cre-dependently expressing ChR2 were injected into the PBel of *Calca*^*Cre/+*^ mice and AAVs Cre-dependently encoding NEP_LDCV_-P2A-mRuby or mCherry were co-injected in the same mice. Four weeks after the injections, the optogenetically evoked excitatory postsynaptic currents (oEPSCs) and potentials (oEPSPs) were recorded in the CeAl neurons that were surrounded by the CGRP^PBel→CeAl^ perisomatic synaptic terminals with mCherry or mRuby fluorescence to validate whether the NEP_LDCV_ selectively silence peptidergic transmission without altering glutamatergic transmissions in CGRP^PBel→CeAl^ synapses (Figure 4E). The oEPSCs were recorded to monitor the glutamatergic transmission, then the oEPSP were recorded in the same neuron to monitor long-lasting resting membrane potential changes induced by trains of 40-Hz stimulation, which is the characteristic response of peptidergic transmission. The whole-cell, patch-clamp recording results revealed that the NEP_LDCV_ had no effect on oEPSCs compared to controls (Figure 4F), whereas it significantly attenuated the oEPSP induced by 40-Hz stimulation in CEAl neurons compared to controls (Figure 4G). These results indicate that the NEP_LDCV_ selectively attenuated the peptidergic transmission leaving the glutamatergic transmission unaltered.

**Figure 4.**
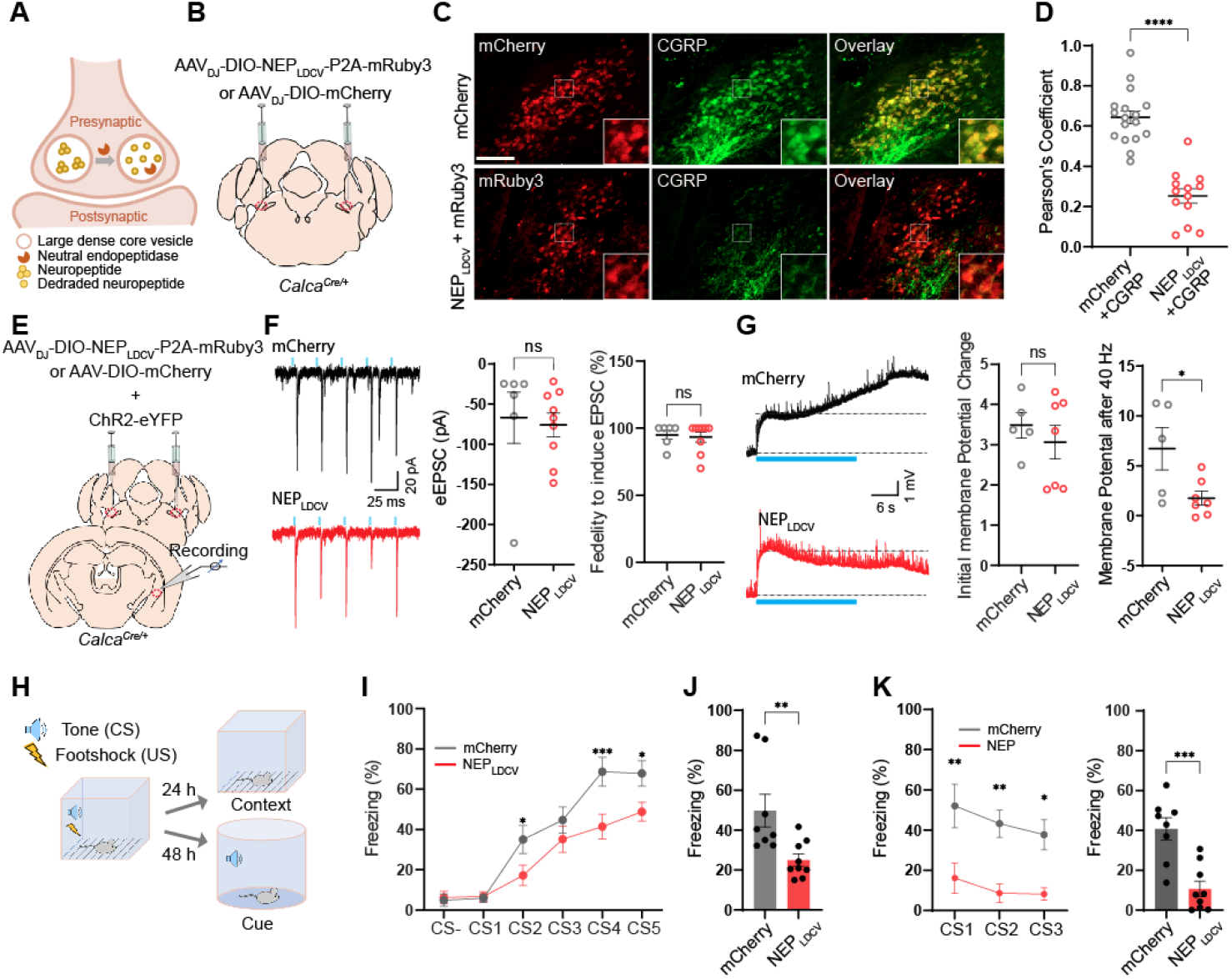
NEP_**LDCV**_ lowers neuropeptide release and attenuates threat learning. (A) Schematic illustrating working principle of LDCV targeted NEP (NEP_LDCV_). (B) Schematic of bilateral stereotaxic injection of NEP_LDCV_ and mCherry into the PBN of *Calca*^*Cre/+*^ mice. Scale bar is 100 μm. (C) Representative images showing mCherry or mRuby3 expressing neurons co-labeling CGRP positive neurons (green) (Scale bar, 100 µm). (D) Quantification of CGRP co-localization in the NEP_LDCV_ sections (n=13 sections form three mice), and mCherry sections (n=17 sections from three mice) by Pearson’s coefficient. ****p < 0.0001 via Two-tailed unpaired t-test comparisons. Data are represented as mean ± SEM. (E) Schematic of viral injection and whole cell recording of *Calca*^*Cre/+*^ PBN slices expressing ChR2 and mCherry or NEP_LDCV_. (F) Example traces (left; Top, ChR2 + mCherry slice. Bottom, ChR2 + NEP_LDCV_ slice.), amplitude (right), and fidelity (right) of oEPSCs in CeA neurons elicited by photostimulation of ChR2-expressing CGRP^PBel^ axonal terminals. n=6 for mCherry, n=9 for NEP_LDCV_. ns, not significant. Data are represented as mean ± SEM. (G) Example traces of oEPSP (left; Top, ChR2 + mCherry slice. Bottom, ChR2 + NEP_LDCV_ slice.), initial oEPSP amplitude (middle), and sustained oEPSP amplitude (right) after 40-Hz photostimulation. ns, not significant. *p < 0.05 via two-tailed unpaired t-test comparisons. Photostimulation onset is indicated by blue line. n=5 for mCherrry, n=7 for NEP_LDCV_. Data are represented as mean ± SEM. (H) Schematic illustration of auditory fear conditioning experiment. (I) Freezing during fear conditioning in mice expressing mCherry (n = 8) and NEP_LDCV_ (n = 9). \**P*<0.05, \*\*\**P*<0.001 via repeated measures two-way ANOVA with Sidak’s multiple comparisons test. Data are represented as mean ± SEM. (J) Freezing at the same context 24 hr after fear conditioning. **p < 0.01 via unpaired t-test comparisons to mCherry. Data are represented as mean ± SEM. (K) Freezing to the tone 48 hr after fear conditioning. Left, \**P*<0.05, \*\**P*<0.01 via repeated measures one-way ANOVA. Right: \*\*\**P*<0.001 via unpaired t-test comparisons. Data are represented as mean ± SEM.

We then investigated whether the degradation of neuropeptides by the NEP_LDCV_ has an impact on Pavlovian threat learning. In the auditory fear conditioning (Figure 4H), we found that the gradual increase of freezing behaviors was observed in the mCherry-expressing control, while NEP_LDCV_-expressing group exhibited a significant reduction of freezing behavior compared to the control group (Figure 4I). In memory tests 24 hr later, mice expressing the NEP_LDCV_ exhibited marked suppression of freezing behaviors in response to the context cue (Figure 4J) and auditory cues (Figure 4K) as compared with the mCherry-expressing group. In addition, we examined whether lowering neuropeptide release by NEP_LDCV_ in CGRP^PBel^ neurons affects animals’ responses to formalin-induced inflammatory pain and quinine, a bitter tastant (Kang et al., 2022). In the formalin assay, formalin-induced licking behaviors were significantly reduced during acute and inflammatory phases (Figure S4A-C). Furthermore, the NEP_LDCV_ group showed significantly increased quinine consumption as compared with mCherry control group (Figure S4D). Overall, these results demonstrated that the NEP_LDCV_ efficiently and selectively lowered neuropeptide release in behaving mice making it an ideal tool to study the direct involvement of peptidergic transmission in the peptidergic circuits.

### Glutamatergic transmission is not involved in threat learning

Since aversive sensory stimuli failed to trigger the SypSEP fluorescence increase at CGRP^PBel→CeAl^ terminals, we investigated whether the glutamate release by CGRP^PBel^ is dispensable in Pavlovian treat learning. To disturb glutamate release, we edited the *Slc17a6* gene, which encodes Vglut2, using the CRISPR/saCas9 system (Hunker et al., 2020), based on a report showing that the Vglut2 is the main glutamate transporter in the PBel (Pauli et al., 2022). AAVs that express Cas9 and sgRNAs for *Slc17a6* or *Rosa26* (as control) in a Cre-dependent fashion were bilaterally injected into the PBel of *Calca*^Cre/+^ mice. Note that *Rosa26* expression has no biological effect since it does not encode a functional protein. Therefore, sg*Rosa26* is used as a control for sg*Slc17a6* (Figure 5A). *In situ* hybridization for *Slc17a6* and *Calca* genes in PBel showed that *Slc17a6* mRNA was depleted in CGRP neurons of sg*Slc17a6*-injected mice but not in sg*Rosa26*-injected mice (Figure 5B).

**Figure 5.**
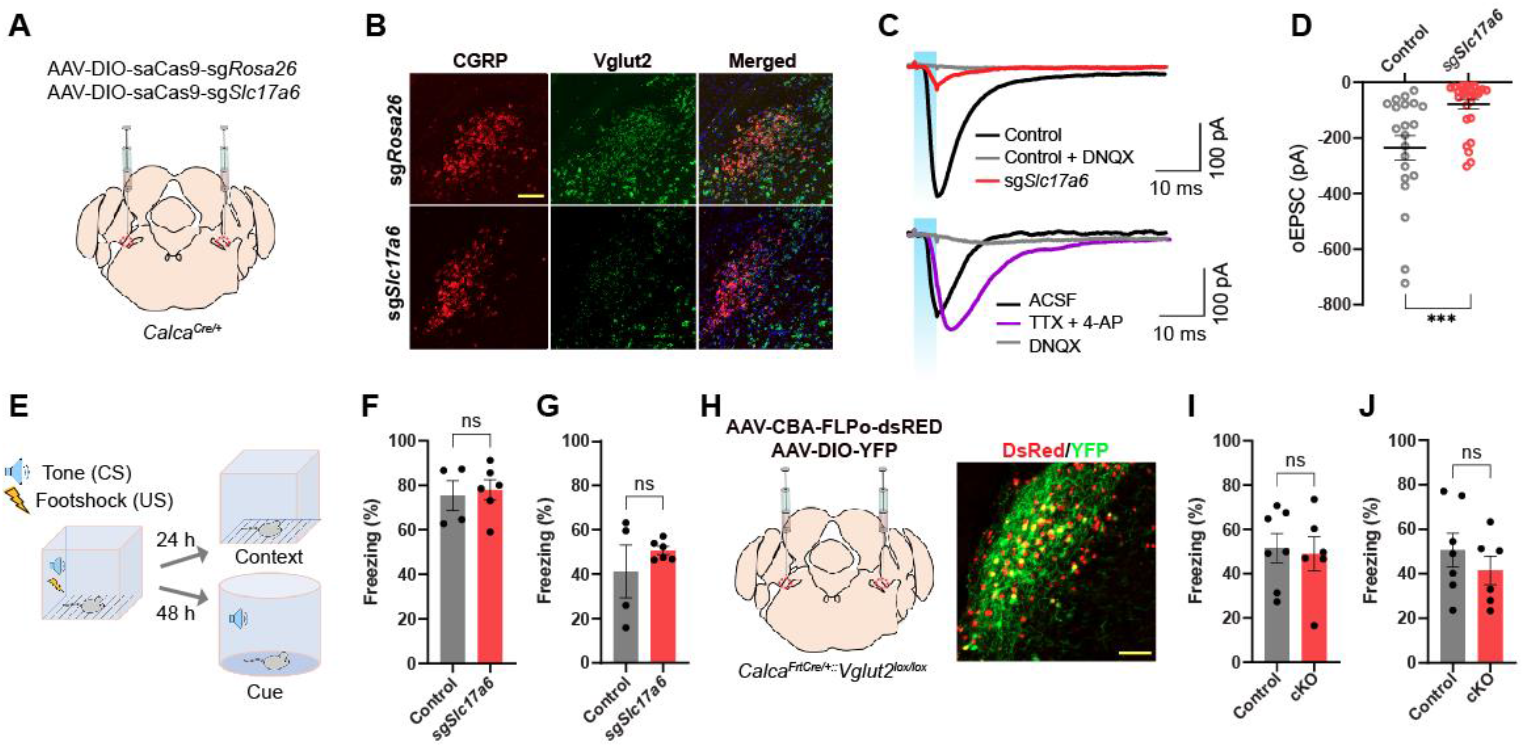
CRISPR and genetic disruption of glutamate release by CGRP^PBel^ does not influence on Pavlovian threat conditioning. (A) Schematic depiction of guide RNAs and saCas9 expression in CGRP^PBel^ neurons. (B) Representative images of *in situ* hybridization detecting *Calca* (encodes CGRP) and *Slc17a6* (encodes Vglut2) in PBL. (C) Example traces of evoked EPSCs in CeA neurons elicited by optogenetic stimulation of axonal terminals of ChR2-expressing CGRP^PBel^ neurons. Top, EPSC traces of control (*sgRosa26*) and *sgSlc17a6* group. Bottom, EPSC traces with the bath application of TTX and 4-AP or DNQX in *sgSlc17a6* group. (D) Amplitudes of EPSCs (n = 21 for control, n = 30 for *sgSlc17a6*). ****p < 0.0001 via Two-tailed unpaired t-test comparisons. Data are presented as mean ± SEM. (E) Freezing during fear conditioning in control (n = 4) and *sgSlc17a6* mice (n = 6). Scale bars are 100 μm. (F and G) Percent of time to spend freezing at the same context 24 hr (F) and to the tone 48 hr (G) after learning in mice expressing sg*Slc17a6* (n = 4) or sg*Rosa26* (n = 6). ns via Two-tailed unpaired t-test comparisons. Data are presented as mean ± SEM. (H) Schematic and histology of *Scl17a6* conditional knockout in CGRP^PBel^ neurons. (I and J) Percent of time to spend freezing at the same context 24 hr (F) and to the tone 48 hr (G) after learning in control (n = 7) and cKO (n = 6) mice. ns via Two-tailed unpaired t-test comparisons. Data are presented as mean ± SEM.

To confirm knockdown of glutamatergic transmission, slice electrophysiology was used to measure oEPSCs in postsynaptic CeAl neurons. AAVs Cre-dependently expressing ChR2 and sg*Slc17a6* were stereotaxically injected into the PBel of *Calca*^Cre/+^ mice. Three weeks after surgery, whole-cell, patch-clamp recording of CeAl neurons showed that sg*Slc17a6* resulted in a significant reduction of oEPSCs compared to controls; responses were mediated by monosynaptic glutamatergic transmission (Figure 5C lower trace). The average amplitude of oEPSCs exhibited by CeAl neurons showed 78.3 % of reduction in sg*Slc17a6*-expressing mice compared to controls. Majority of neurons (21/30) did not show any oEPSCs and 30 % of remaining neurons showed reduced oEPSCs (Figure 5C upper trace, and 5D). We then performed Pavlovian threat conditioning with sg*Slc17a6* and control groups (Figure 5E). Animals in both groups showed a gradual increase in freezing behavior as the number of pairings increased (Figure S5A). Furthermore, both groups of mice showed similar levels of freezing in contextual (Figure 5F) and auditory cue-induced (Figure 5G) threat memory tests.

To further test the contribution of glutamatergic transmission in threat learning, *Slc17a6* conditional knockout (cKO) mice were produced by crossing the *Slc17a6* floxed mice (*Slc17a6*^*l*ox/lox^) with mice that express Cre-recombinase specifically in CGRP neurons in a Flp-dependent manner (*Calca*^*Frt*Cre/+^). Bilateral injection of AAV encoding Flp-recombinase into the PBel of *Calca*^*Frt*Cre/+^::*Slc17a6*^lox/lox^ mice induced conditional knockout of the *Slc17a6* gene specifically in CGRP^PBel^ neurons (Figure 5H). Both cKO and control mice displayed similar freezing responses during threat learning (Figure S5B), as well as contextual-(Figure 5I) and cue-dependent retrieval tests (Figure 5J). These results show that lowering glutamate release from CGRP^PBel^ neurons had no effect on threat learning.

## DISCUSSION

Here we report the development of two new genetically encoded tools that can monitor and silence the release neuropeptides in awake behaving mice: The CybSEP2, a genetically encoded fluorescent LDCV sensor that reliably detects LDCV release presynaptically in freely moving mice, and the NEP_LDCV_, a genetically encoded peptidase that attenuates neuropeptide release by degrading neuropeptides inside the LDCVs in freely moving mice.

Recent progress in developing several neuropeptide sensors that utilize postsynaptic neuropeptide receptors now allows one to monitor the release of some neuropeptides in behaving animals. These technologies will provide important insights into the function of neuropeptides in brain circuits (Sabatini and Tian, 2020; Wu et al., 2022), but they have some limitations. First, the GPCR-based sensor is expressed in postsynaptic neurons and detects the release of neuropeptides indirectly, regardless of their source. In cases where a postsynaptic neuron receives peptidergic inputs from different brain areas, it is not possible to distinguish which synaptic terminal evoked the release event using a postsynaptic sensor. In addition, it has been suggested that neuropeptides can be released from anywhere in a neuron, including the cell body, dendrites, and axonal terminals (Ludwig and Leng, 2006; van den Pol, 2012). Thus, GPCR-based sensors cannot pinpoint the exact neuropeptide release site. Second, the receptors for some neuropeptides, such as cocaine and amphetamine-related transcript (CART), have not yet been discovered (Ahmadian-Moghadam et al., 2018). Thus, GPCR-based sensors cannot be applied to these neuropeptides. Third, the GPCR-based sensors are artificially expressed in postsynaptic neurons at high level, and therefore can act as competitive inhibitors of the endogenous GPCR by capturing endogenous ligands even if their downstream signaling actions are eliminated by genetic modification. This could potentially cause abnormal behavior in animals. Finally, one GPCR-based sensor usually detects the release of only one neuropeptide, which provides high sensitivity for a specific neuropeptide release event but limits its versatility. Considering more than a hundred neuropeptides and their postsynaptic GPCRs have been discovered, more than a hundred individually developed GPCR-based sensors are needed. These limitations would be resolved by developing a presynaptic neuropeptide release sensor. Our CybSEP2 system: 1) can pinpoint exactly where the neuropeptides are released from, 2) can be used to monitor essentially any neuropeptide by driving the expression of CybSEP2 via a neuropeptide-specific Cre-driver mouse line, and 3) does not interfere with endogenous peptidergic signaling. Given the fact that only a handful of GPCR-based neuropeptide sensors are currently available (Dong et al., 2022; Qian et al., 2022; Wang et al., 2022), CybSEP2 can serve as an alternative tool for monitoring the release of neuropeptides for which postsynaptic sensors are not yet available. However, this versatility limits the specificity of the CybSEP2 system. Considering that multiple neuropeptides can be co-packaged into the same LDCV, monitoring LDCV release events as a proxy for neuropeptide release cannot define which neuropeptide has been released. Therefore, current pre- and post-synaptic neuropeptide sensors each have limitations, but when used in combination their unique properties could lead to transformative discoveries.

Most peptidergic neurons are multi-transmitter neurons (Hokfelt, 1991; Lundberg, 1996; Merighi, 2002). Therefore, to tease apart the roles played by each transmitter in single neurons, a silencing tool that selectively inhibits the release of specific transmitter in a defined peptidergic circuit is critically required, but such a tool is currently unavailable. Genetic knock-out of the neuropeptide encoding gene is one way to interrogate the role of certain neuropeptide. Yet, considering most peptidergic neurons co-express and co-package multiple neuropeptides in a single LDCV, knocking-out one neuropeptide gene may be insufficient to produce phenotypic changes due to the compensatory actions of other neuropeptides co-expressed in the same neurons (Zajdel et al., 2021). Therefore, developing a tool that silences the release of all neuropeptides in a genetically defined neuronal population is required to interrogate peptidergic transmission systemically. In this study, we selectively silenced the peptidergic transmission by targeting the NEP into the luminal side of the LDCVs that proteolytically degrade neuropeptides inside the LDCVs leaving fast synaptic transmission unaltered. The NEP, a zinc-dependent metalloprotease, is an integral plasma membrane protein that primarily degrades neuropeptides from the extracellular surface (Booth and Kenny, 1980). The enzymatic activity of the NEP at pH 5.0 is comparable to its activity at pH7.4 (Kerr and Kenny, 1974). Therefore, the LDCV-targeted NEP (NEP_LDCV_) can reliably and selectively degrade neuropeptides at low pH in the LDCV. Our results showed that the cell-type-specific expression of the NEP_LDCV_ in the CGRP^PBel→CeA^ peptidergic circuit substantially reduced peptidergic transmission without altering glutamatergic transmission by selectively degrading neuropeptides in the CGRP^PBel^ neurons, which resulted in impaired responses to innately aversive sensory stimuli, such as electric footshock, quinine consumption, and the plantar injection of formalin (Figure 4). These results indicate that the AAV-DIO-NEP_LDCV_-P2A-mRuby3 can be used as the specific silencer for the transmission of many neuropeptides that contain hydrophobic amino acid in awake behaving animals.

With the novel sensor and silencer for neuropeptide release at hand, we asked whether neuropeptides can function as the primary transmitter in mediating a major output, or whether they invariably act as a co-transmitter to modulate classical neurotransmission. Pharmacological blockade of postsynaptic neuropeptide receptors substantially affected neuronal outputs and behavior, indicating that peptidergic transmissions play major roles in certain peptidergic systems in the brain (Hokfelt et al., 2003; Salio et al., 2006). Yet, it remained unclear whether neuropeptides can be the only released transmitter or whether they are always co-released with other transmitters to affect behavior or physiology in mammals (Salio et al., 2006). To address this question, we monitored the release of two types of transmitter vesicles, LDCVs and SVs, using our newly developed CybSEP2 together with a previously developed SV sensor, SypSEP, during threat learning. We investigated the CGRP^PBel→CeA^ peptidergic circuit since our previous analyses have shown that CGRP^PBel^ neurons and their direct downstream targets (neurons in the CeA that express the CGRP receptor) are critically involved in affective pain transmission and aversive memory formation (Han et al., 2015). Using a CMOS-coupled fiber photometry system, we observed that noxious stimuli such as an electric footshock, exposure to a hot plate, or quinine consumption triggered the release of neuropeptides including CGRP in the CGRP^PBel→CeA^ peptidergic pathway, but not glutamate (Fig. 3). Interestingly, noxious, and aversive taste stimuli decreased SypSEP fluorescence below the baseline in the same circuit (Fig. S3). These results suggest that only neuropeptides are released from CGRP^PBel→CeA^ terminals evoked by aversive sensory stimuli. Furthermore, functional silencing of peptidergic or glutamatergic transmissions in this pathway using the NEP_LDCV_, or the *Slc16a7* gene disruption further confirmed that peptidergic, but not glutamatergic transmission is required for conveying aversive US from the PBel to the amygdala during Pavlovian threat learning. It is surprising that glutamatergic transmission is dispensable for relaying aversive sensory cues during threat learning. Previous studies showed that glutamatergic transmission in the CGRP^PBel→CeA^ peptidergic pathway mediate hypercapnic arousal and sleep regulation (Kaur et al., 2013; Kaur et al., 2017). These results suggest that different transmitters in the same neural circuit may play different roles, which raises an intriguing question whether single CGRP^PBel^ neurons can encode distinct information by releasing different transmitters, or two distinct CGRP^PBel^ subpopulations exclusively use different transmitters to mediate distinct functions. Furter investigation is required to address this question.

Taken together, our results show that the CybSEP2 sensor and the NEP_LDCV_ silencer are reliable tools for monitoring and silencing presynaptic neuropeptide release in awake behaving mice as they experience sensory or emotional stimuli. Furthermore, these new tools will allow researchers to investigate molecular and cellular mechanisms of peptidergic transmission more thoroughly, which has been largely neglected compared to the fast synaptic transmission.

## Acknowledgments

We thank Dr. David O’Keefe and Han lab members for critical discussion of this manuscript. We also thank Susan Phelps for animal husbandry and James Allen for helping with custom AAV production. S.H. is supported by 5R01MH116203 from NIMH, 1RF1NS128680 from NINDS, and the Salk Institute Innovation Grant.

## Author Contributions

S.H. conceived of the idea. S.H., R.D.P and L.S.Z. secured funding. S.H., D-I.K., S.P., and J.Y.C. designed the experiments and wrote the manuscript. D-I.K. cloned the sensor and silencer of peptidergic transmission and performed most of the experiments. S.P. performed the *Slc17a6* loss-of-function experiments. M.Y. and J.Y.C. performed the slice electrophysiology associated with the NEP_LDCV_, and *Slc17a6* loss-of-function experiments. J.J. built the custom-made CMOS fiber photometry system. A.C.H, and L.S.Z. provided AAVs Cre-dependently expressing Cas9 and sgRNAs for *Slc17a6* or *Rosa26*.

## STAR METHODS

### EXPERIMENTAL MODEL AND SUBJECT DETAILS

#### Mouse lines

All protocols for animal experiments were approved by the IACUC of the Salk Institute for Biological Studies and University of Washington according to NIH guidelines for animal experimentation. *Calca*^*cre*^, *Slc17a6*^*lox/lox*^, and *Calca*^*FrtCre*^ transgenic mouse lines used in this study are all C57BL/6J background, and generated from the Palmiter lab (Hnasko et al., 2010; Carter et al., 2013; Chen et al., 2018). 3–4-month-old heterozygous mice were used for all experiments except for the *Slc17a6* experiments where homozygous for *Slc17a6*^*lox/lox*^ were used. Animals were randomized to experimental groups, and no sex differences were noted. Mice were maintained on a standard 12-hour light/dark cycle and provided with food and water *ad libitum*.

## METHOD DETAILS

### Construct design and molecular cloning

To generate fusion constructs, cytochrome b561-superecliptic pHluorin (CybSEP), mouse cytochrome b561 (NM_007805.4, Genscript) and pcDNA3-SypHluorin2 (#37005, Addgene) were used for this construct. We first created point mutated cytochrome b561 by substituting two extracellular histidines (His 86 and 159) by two alanines, which are a critical role in binding of ascorbic acid. Then, pHluorin was inserted into the lumen domain at position 339-340 between transmembrane domain 3 and 4 of cytochrome b561. For two copies of pHluorin, the second one was subsequently inserted into the backbone vector. For cloning cytochrome b561-gamillus construction (124837, Addgene), pHluorin was replaced with gamillus. For adeno-associated virus (AAV) constructs, we used rAAV-hSyn backbone (51509, Addgene) and *Asc*I and *Fse*I were used to excise the insert of the backbone for replacement of fusion constructs. For LDCV targeted peptidase (NEP_LDCV_) generation, the signal peptide (0-26 amino acids) of pro-opiomelanocortin (#176704, Addgene) and ectodomain (52-750 amino acids) of the NEP (#7283, Addgene) were used for the fusion protein and then inserted in rAAA-FLEX-axonGCaMP6s-P2A-mRUBY3 vector (#112008, Addgene) used as a backbone after excising axonGCaMP6s by *Bam*HΙ and *Nhe*Ι. All construct generation was performed by In-Fusion HD cloning kit (638920, Takara).

### Cell culture and live-cell imaging

PC-12 cells (CRL-1721, ATCC) were maintained in DMEM high glucose (#11995065, Invitrogen) supplemented with 10% fetal bovine serum (#10437028, Hyclone) and 5% horse serum (<u>16050130</u>,GIBCO) at 37°C incubator with 5% CO_2_. For imaging, cells were plated on the poly-L-lysine (Sigma) coated coverslips in 24-well plates and following day plasmids were transfected using by lipofectamin 3000 (#L3000015, Invitrogen) and experiments were performed 48 hr after transfection. To differentiate cells, 50 ng/ml nerve growth factor (NC010, Sigma) was added to the plates 12 hours after transfection. For pH-dependent experiments, CybSEP or CybGam-expressing cells were perfused with extracellular bath solution and with acidic extracellular solution at pH5.5 to quench the pHluorin. 50 mM NH4Cl was then added to the chamber to neutralize the cellular environment. To elicit the membrane fusion of large dense core vesicles by depolarization, 70 mM KCl was added on the cells grown on coverslips. For the electrical stimulation, broken glass pipettes were pulled from borosilicate glass (G150TF-4, Warner Instruments) with pipette puller (SU-P97, Shutter instrument) and filled with the extracellular bath solution. For stimulation, they were positioned near fluorescence expressing cells and were evoked by electrical stimulator (Model 2100 Isolated Pulse Stimulator, A-M systems). The stimulation voltage was set at 4-5 V (3-ms pulse width) with various frequencies to elicit release of neuropeptides from LDCVs. During perfusion, the temperature of the bath chamber was maintained at 32°C by in-line solution heater (TC-324C, Warne Instruments). The extracellular bath solution contained (in mM) 130 NaCl, 2.8 KCl, 2 CaCl2, 1 MgCl_2_, 10 HEPES and 10 glucose; for cellular acidification the acid solution contained (in mM) 90 NaCl, 2 CaCl_2_, 1 MgCl_2_, 60 Na-acetate and 10 HEPES. For the cellular neutralization, the 40 mM NaCl of bath solution was replaced with 40 mM NH_4_Cl. To test the role of calcium in the release of neuropeptides, 5 mM EGTA was replaced with CaCl_2_.

Imaging was carried out in upright scope (Slice scope Scientifica) with water immersion objective lens (5X, 0.1 NA; 40X, 0.9 NA, LUMPLFLN-W, Olympus), LED illumination (GFP, 490-nm; mCherry, 580-nm) (pE-4000, CoolLED) and a multiband filter set (89402, Chroma). Images were acquired at 10 Hz (2 × 2 digital binning, 1,024 × 1,024-pixel resolution) equipped with an sCMOS camera (Prime 95B, Teledyne Photometrics) using micro-manager open-source software. Quantification and statics analysis image data from cultured cells and brain slices were processed with Image J or Prism (Graphad). ROIs were manually selected by live scanning function of ImageJ after photobleach correction. The fluorescent signal from each cell was obtained by averaging fluorescent changes (ΔF/F_0_) of individual cells. The fluorescence responses (ΔF/F_0_) were calculated as (F_raw_ –F_baseline_)/F_baseline_.

### AAV viral preparation

All AAV viruses used in the study were generated in the lab as previously described with a minor modification except AAV-DIO-mCherry (#50459, Addgene), AAV_DJ_-EF1a-DIO-hChR2(H134R)-EYFP-WPRE-pA (# 20298, Addgene), AAV1-DIO-saCas9-sg*Slc17a6* (The Zweifel Lab), AAV1-DIO-saCas9-sg*Rosa26* (The Zweifel Lab), AAV1-CBA-FLPo-dsRed (The Palmiter Lab) and AAV1-hSyn-DIO-YFP (The Palmiter Lab). In brief, constructs cloned in AAV-hSyn vectors were transfected with RC/DJ and adenovirus-helper plasmid into the AAV-293 cells (#240073, Agilent) using calcium phosphate precipitation method. 72 hr after transfection, cells were harvested, lysed, and collected by centrifugation to remove debris. AAV particles were subsequently purified using a HiTrap heparin column (GE healthcare, UK) and concentrated by Amicon ultra-4 centrifugal filter (UFC801008, MilliporeSigma).

### Stereotaxic surgery

All surgeries were carried out under 1.5%-2% isoflurane anesthesia (Dräger Vapor® 2000; Draegar) and kept on the water circulating heating pad to maintain body temperature during surgery. Mice were place in a stereotaxic frame (David Kopf Instruments). Viral Injections were unilaterally or bilaterally delivered using a syringe (65458-01, Hamilton, USA) controlled by an ultra-micropump (UMP-3, World Precision Instruments, USA) at a rate of 0.1 µl/min (total volume of 0.6 µl for slice imaging and 0.4 µl for fiber photometry). For slice imaging, AAVDJ-hSyn-DIO-CybSEP, AAV_DJ_-hSyn-DIO-CybGam, AAVDJ-hSyn-DIO-mCherry, or/and AAV_DJ_-hSyn-DIO-TetTox-mCherry were bilaterally injected into the PBN of *Calca*^*Cre/+*^ (AP: −5.2 mm, ML: ±1.5 mm, DV: −3.6 mm). For fiber photometry recording, AAV_DJ_-hSyn-DIO-CybSEP or AAV_DJ_-hSyn-DIO-CybGam was unilaterally injected into the PBN of *Calca*^*Cre/+*^ and a custom-made optic ferrule (0.4 uM, 0.5 NA) was implanted into the CeA of *Calca*^*Cre/+*^ (AP: −1.2 mm, ML: −2.85 mm, DV: −4.5 mm). Optic ferrule was then implanted above the injection site (AP: −1.2 mm, ML: −2.85 mm, DV: −4.5 mm). Mice were allowed to recover for at least 3 weeks following viral infection and fiber implantation before behavioral testing. For NEP_LDCV_ expression, AAV_DJ_-hSyn-DIO-NEP_LDCV_-P2A-mRuby2 or AAV_DJ_-hSyn-DIO-mCherry were injected bilaterally into the PBN of *Calca*^*Cre/+*^. For patch clamp, AAV_DJ_-hSyn-DIO-NEP_LDCV_-P2A-mRuby2 or AAV_DJ_-hSyn-DIO-mCherry with AAV1-DIO-ChR2-YFP were bilaterally injected into the PBN of *Calca*^*Cre/+*^. For *Slc17a6* gene inactivation, AAV1-DIO-saCas9-sg*Slc17a6* or AAV1-DIO-saCas9-sg*Rosa26* were bilaterally injected into the PBel (AP, −4.9 mm; ML, ±1.35 mm; DV, 3.5 mm) at a rate of 0.1 μl/min (total 0.5 μl). For For patch clamp, AAV1-DIO-saCas9-sg*Slc17a6* and AAV1-DIO-ChR2-YFP were bilaterally injected into the PBel.

### Histology and imaging

Mice were euthanized with CO2 at a flow rate of 3 L/min (LPM) and transcardially perfused with cold paraformaldehyde (PFA) (4% in PB). Brains were kept in PFA overnight at 4 °C and dehydrated in 30% sucrose (in PBS) for at least 18 hours before vibratome sectioning (VT1000s, Leica). Brains were cut into 50 µm coronal section using cryostat (CM 1950, Leica), collected in PBS and mounted on Superfrost microscopic slides (Fisher Scientific) with DAPI Fluoromount-G mounting media (Southern Biotech) for imaging. For immunostaining with CGRP antibody, coronal slices were obtained from AAV_DJ_-hSyn-DIO-NEP_LDCV_-P2A-mRuby2 or AAV_DJ_-hSyn-DIO-mCherry expressing mice. Slices then were washed with PBST containing 0.1% Tween-20s and blocked with 3% normal donkey serum for 1 hour at room temperature. After rinsing with PBST, slices were incubated with rabbit anti-CGRP (1;1000) at 4 °C overnight. Next day, slices were rinsed with PBST and then incubated with anti-rabbit Alexa Fluor® 488-secondary antibody for 1 hour. Images of slices were acquired using all-in-one fluorescence microscope (BZ-X710, Keyence) with objective lens (10X, 0.40 NA; 20X, 0.75 NA, Olympus) with the BZ-X viewer software.

### Fluorescence imaging in acute brain slices

Three weeks after viral injection into PBN, acute brain slices containing PBN (200 µm) and CeA (250 µm) were prepared for imaging. Mice were deeply anesthetized by isoflurane prior to decapitation and transcardial perfusions were performed with 50 mL of ice-cold carbogenated (95% O2 :5% CO2) cutting solution (110.0 mM choline chloride, 25.0 mM NaHCO3, 1.25 mM NaH2PO4, 2.5 mM KCl, 0.5 mM CaCl2, 7.0mM MgCl2, 25.0 mM glucose, 5.0 mM ascorbic acid and 3.0 mM pyruvic acid). Brains were immediately removed and mounted on the chamber of a VT 1200S Vibratome (Leica) followed by 200 µm thick sections by vibratome in the same solution. Slices were transferred recovery chamber containing carbogenated artificial cerebrospinal fluid (aCSF; 124 mM NaCl, 2.5 mM KCl, 26.2 mM NaHCO3, 1.2 mM NaH2PO4, 13 mM glucose, 2 mM MgSO4 and 2 mM CaCl2). After recovery at 34°C water bath for 15 min, slices were transferred to room temperature for at least 30 min before imaging and then slices were transferred to the imaging chamber and perfused with carbogenated aCSF solution with the flow rate of 2 ml/ min and the temperature of the chamber was maintained at 30–34°C by a temperature controller (TC-324C, Warne Instruments). For fluorescence imaging in cell bodies or axonal terminals, the electrical stimulation and acquisition were applied the same condition as in the cultured cells.

### Slice electrophysiology

Mice were anesthetized with pentobarbital sodium and phenytoin sodium solution (Euthasol, 0.2 ml, i.p.) and transcardially perfused with ice-cold cutting solution (92 mM N-methyl-D-glucamine, 2.5 mM KCl, 1.25 mM NaH2PO_4_, 30 mM NaHCO_3_, 20 mM HEPES, 25 mM D-glucose, 2 mM thiourea, 5 mM Na-ascorbate, 3 mM Na-pyruvate, 0.5 mM CaCl_2_, 10 mM MgSO_4_). Mice were decapitated, brains quickly removed and chilled in ice-cold cutting solution. Coronal slices (300 µm) were cut with a vibratome (Leica VT1200) and incubating in the same cutting solution at 33°C for 12 min. Slices were transferred to a storage chamber containing recovery solution at room temperature (124 mM NaCl, 2.5 mM KCl, 1.25 mM NaH2PO4, 24 mM NaHCO3, 5 mM HEPES, 13 mM D-glucose, 2 mM CaCl2, 2 mM MgSO4). Slices were transferred into the recording chamber maintained at 33 °C and perfused with artificial cerebral spinal fluid (126 NaCl, 2.5 KCl, 1.2 NaH2PO4, 26 NaHCO3, 11 D-glucose, 2.4 CaCl2, 1.2 MgCl2). Blockers were added to the recording solution for a final concentration of: 10 µM DNQX, 1 µM TTX, and 50 µM 4-AP (Tocris Bioscience). All solutions were continuously bubbled with 95% O2-5% CO2 with pH 7.3-7.4 and 300-310 mOsm. Whole cell patch-clamp recordings were obtained using an amplifier (Molecular Devices, MultiClamp 700B) and filtered at 2 kHz. Patch electrodes (3-5 MΩ) were filled with a cesium-methanesulfonate internal solution (117 mM Cs-methanesulfonate, 20 mM HEPES, 0.4 mM EGTA, 2.8 mM NaCl, 5 mM TEA, 5 mM ATP, 0.5 mM GTP; pH 7.35, 280 mOsm). Only cells with access resistance <20 MOhms throughout the recording were used in the analysis. Peak amplitudes of evoked responses were calculated with the average of 10 traces using Clampfit in the pClamp 11 software suite (Molecular Devices).

### Fiber photometry recording

CybSEP2 response in axonal terminals were recorded through bundle-imaging fiber photometry system equipped with CMOS sensor (Doric lenses) and acquired by a LabVIEW (National Instrument) based platform. To measure fluorescence signals, 470-nm LED was used for inducing Ca^2+^ dependent fluorescence signals and 405-nm LED was used for Ca^2+^ independent (isosbestic control) fluorescence signals at a sampling rate of 10 Hz. Both LEDs were bandpass filtered and passed through a 20X/0.4 NA objective lens (Olympus) coupled with optic ferrule implanted on mice by a custom patchcord (400 µm, 0.48 NA). Photometry data were analyzed using Python script. 405-nm channel (F_405fit_) was fit with least mean squares that was scaled to the 470-nm channel (F_470_). Motion corrected ΔF/F was calculated as ΔF/F = (F_470_ -F_405fit_) / F_405fit_. ΔF/F was then z-scored relative to the mean and SD of the fluorescence signal and smoothed by 2nd-order Savitzky–Golay filter using Prism 8 (GraphPad software).

### Behavioral experiments

#### Hot plate test

For recording fluorescent activity in response to noxious thermal stimulus, mice were tethered to a patch cord and placed inside a transparent Plexiglas cylinder (D = 11 cm, H = 15 cm). The bottom of the cylinder was wrapped by thin foil to facilitate the cylinder transfer with minimizing movement effect. Mice were allowed to freely move in the cylinder at room temperature (RT) for 30 min before measurement. Then the baseline was measured at RT and cylinder was transferred to 42°C or 52°C hot plate (PE34, IITC Life Science) to measure the activity of CybSEP2 during noxious thermal stimulus (cutoff time of 20 s).

### Quinine-induced taste aversion test

For recording fluorescent activity in response to aversive taste stimulus, mice were tethered to a patch cord and placed inside a plastic cylinder (11-cm diameter, 15 cm height) with 2-cm diameter hole to deliver water or 0.5 mM quinine. For the first 2 days, mice were habituated in the cylinder. The day before measurement, mice were water deprived at the home cage overnight. The following day, mice were placed to the cylinder for 30 min prior to recording and then fluorescent signal of the sensors was recorded during licking water or quinine. In the test with NEP_LDCV_ expressing mice, they consumed quinine for 10 min after overnight water deprivation and amount consumed was calculated. For taste preference test, a two-bottle choice test between water and quinine was performed.

### Foot shock fear conditioning

A fear conditioning chamber (26 × 30 × 33 cm, ENV-007CT, MED Associates) consisted of a metal grid floor (ENV-005, MED Associates) and standalone aversive electric shock stimulator (ENV-414S, MED Associates) was employed for footshock (unconditioned stimulus: US) fear conditioning. To deliver a tone (conditioned stimuli; CS+), two speakers (AX210, Dell) were placed right next to the foot shock chamber. On day 1, mice were tethered to the patch cord and habituated inside the foot-shock chamber, which involved six CS+ (30-sec, 2-kHz pure tone) with random inter-event intervals. On day 2, mice were placed to the same chamber and underwent footshock fear conditioning, which involved five CS+ that co-terminated with footshock (0.2 mA, 2 sec) over random inter-event intervals. For contextual fear conditioning on the third day, mice were placed to the foot-shock chamber for 3 min. For the cue test, mice were placed to the new context (a glass cylinder wrapped with a non-transparent material; 20-cm diameter, 15-cm height) and then CS+ was delivered three times without US. EthoVision XT 12 software (Noldus) with GigE USB camera (Imagine Source) was used for video recording, foot shock delivery, and analysis of freezing behavior.

For the fear conditioning experiments in Figure 5 and S5, mice were habituated to the context (28×28×25 cm chamber with metal walls and electric grid bottom, MedAssociate) and a 30 s CS (10 kHz, 70 dB tone) on Day 1. After 2 min of baseline, 6 CS were introduced with randomized intervals (60 s – 180 s). The next day, they were exposed to the same context and the same number of tones co-terminated with foot shock (0.3 mA, 0.5 s). On Day 3, mice were exposed to the same context for 5 min to test context-dependent fear memory. The cue-dependent fear memory test was conducted in a different context (28×28×25 cm chamber with white acrylic panel walls). Mice placed in the different context received 3 CS following of 2 min baseline. Freezing behavior was measured using Ethovision XT 15 software (Noldus). Freezing was determined as the time when the velocity of center point of mouse was under 0.75 cm/s. Freezing levels during the 30 s of CS were counted for day 1 and 3. For the contextual fear memory test, freezing time was measured for last 3 min of the session. The average of 3 CS was calculated to obtain the freezing level for the cue-induced fear memory test.

---

**Figure S1.**
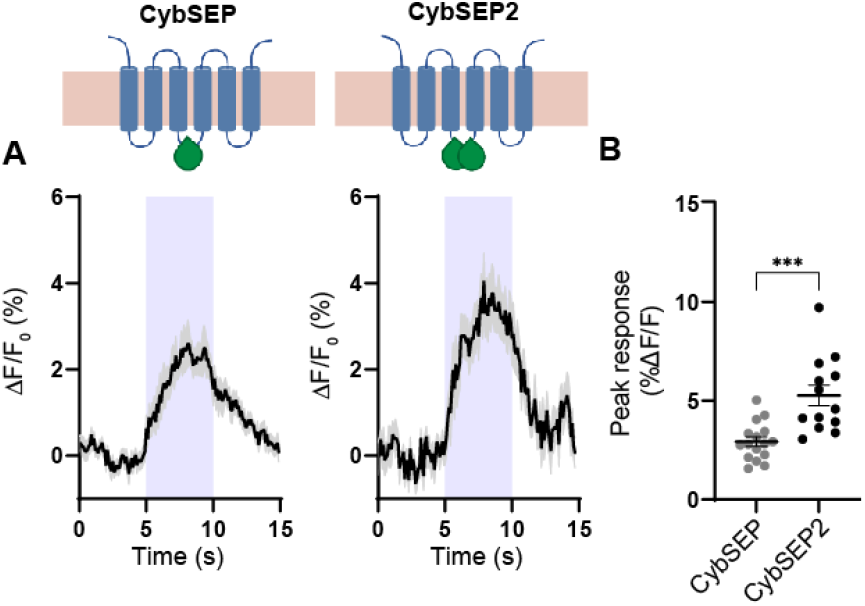
Comparison of fluorescence levels of CybSEP and CybSEP2 evoked by electrical stimulation. (A and B) Schematic and average traces showing comparison between CybSEP (n = 18) and CybSEP2 (n = 13) expressing cells (duplicated from Figures 1G) during electrical stimulation at 100 Hz and quantification of percent ΔF/F_0_ peak intensity in (A) (***p < 0.001 via unpaired t test). Data are represented as mean ± SEM.

**Figure S2.**
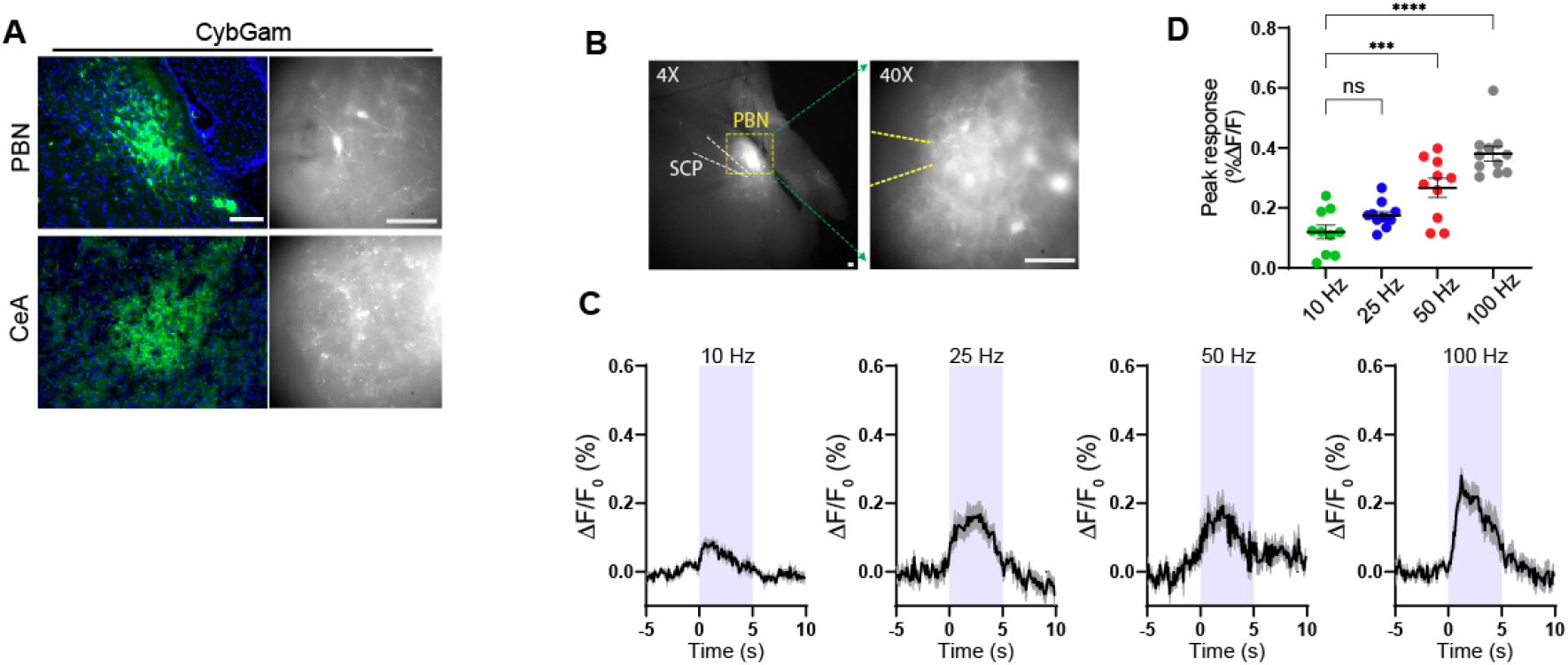
Expression of CybGam and imaging CybSEP2 in acute brain slice containing the PBN. (A) Schematic and images showing expression of CybGam in the PBN and the CeA of *Calca*^*Cre/+*^. (B) Images of PBN showing CybSEP2 expression for slice imaging. SCP indicates superior cerebellar peduncle (Scale bar, 100 µm). (C and D) Average traces of fluorescence change in response to various electrical stimulation and quantification of data in (C) (10-12 traces from 18 slices prepared from 3 mice; ***p < 0.001, ****p < 0.0001 via one way ANOVA followed by Tukey’s multiple comparisons to the 10 Hz). Data are represented as mean ± SEM.

**Figure S3.**
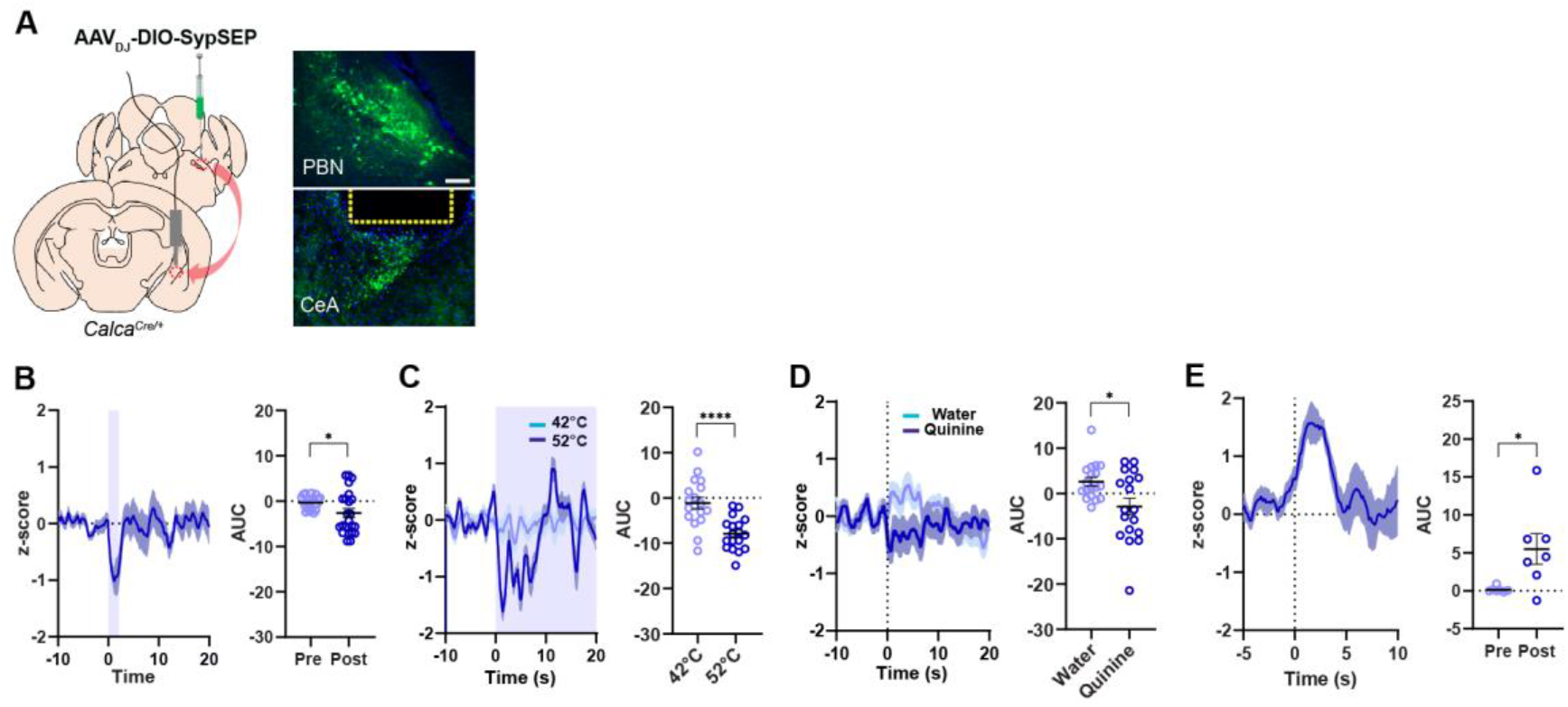
Deep brain recording of the SV sensor in the synaptic terminals of freely moving mice. (A) Schematic illustration of viral injection and images showing expression of SypSEP in the PBN and in the CeA of *Calca*^*Cre/+*^. Yellow dot line represents the location of optic fiber (Scale bar, 100 µm). (B) Average trace of fluorescence change on footshock and quantification of data 10 s before and after footshock (23 traces from 4 mice; *p < 0.05 via paired t test). (C) Average traces of fluorescence change during thermal stimulus and quantification of data for 0-10 s (18 traces from 4 mice; ****p < 0.0001 via unpaired t test). (D) Average traces of fluorescence change during quinine intake and quantification of data for 0-10 s (18 traces from 4 mice *p < 0.05 via unpaired t test). Data are represented as mean ± SEM. (E) Average traces of fluorescence change during Ensure intake and quantification of data for 0-10 s (7 traces from 3 mice *p < 0.05 via unpaired t test). Data are represented as mean ± SEM.

**Figure S4.**
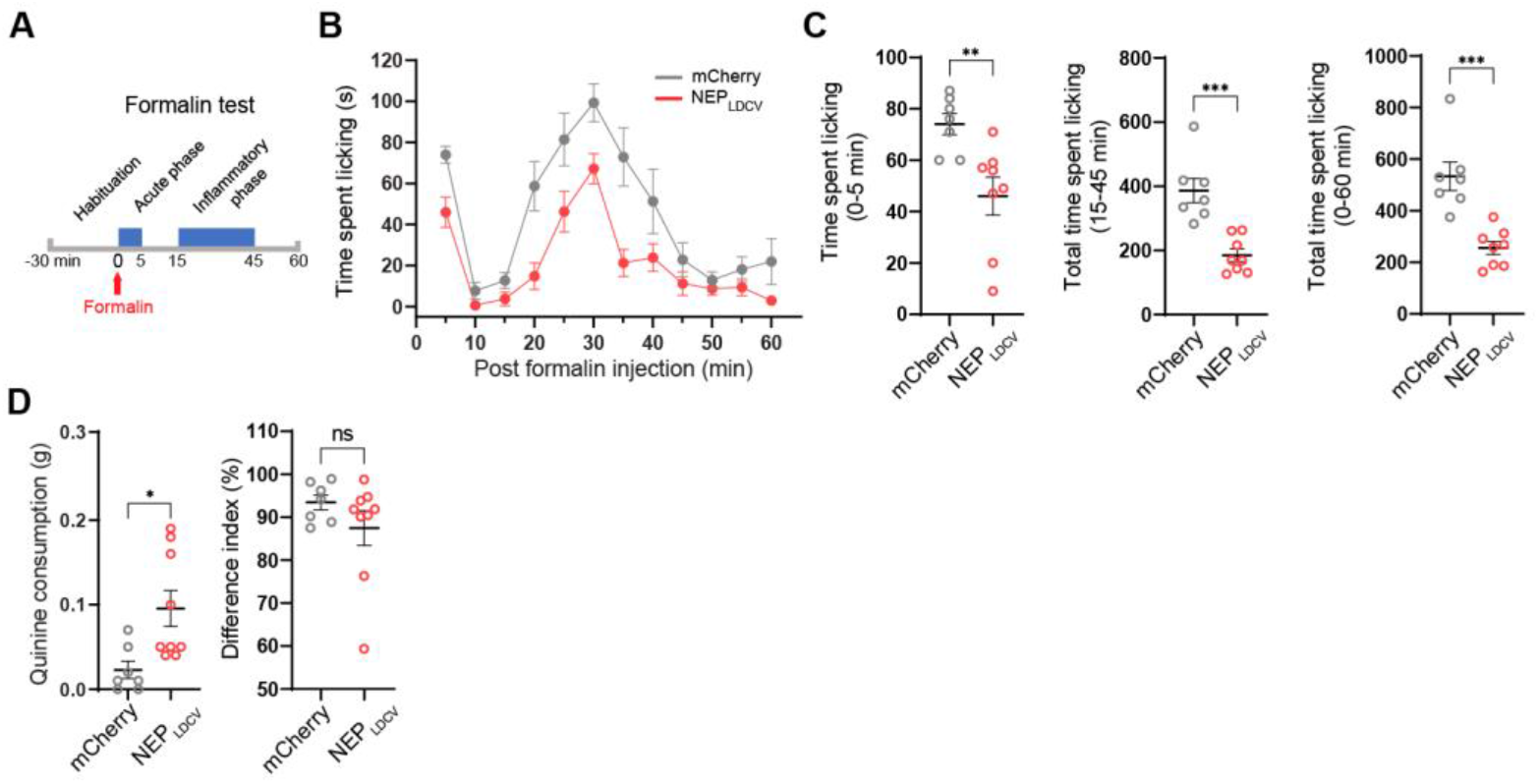
The effect of the NEP_**LDCV**_ on pain behavior by sensory stimulus and formalin injection. (A) Schematic of formalin assay for acute and inflammatory pain tests. (B) Time course of formalin-induced nociceptive responses in the mice expressing mCherry (n=7 for mCherry, n=9 for NEP_LDCV_). (C) Quantification of acute phase (0-5 min, left), inflammatory phase (15-45 min, middle), and a total spent time for locking (0-60 min). \*\**P*<0.01, \*\*\**P*<0.001 via unpaired t-test comparisons to mCherry. Data are represented as mean ± SEM. (D) Quinine consumption (left) and two bottle (right) tests. in the mice expressing mCherry (n = 7) and NEP_LDCV_ (n = 9). \**P*<0.05, ns via unpaired t-test comparisons to mCherry group.

**Figure S5.**
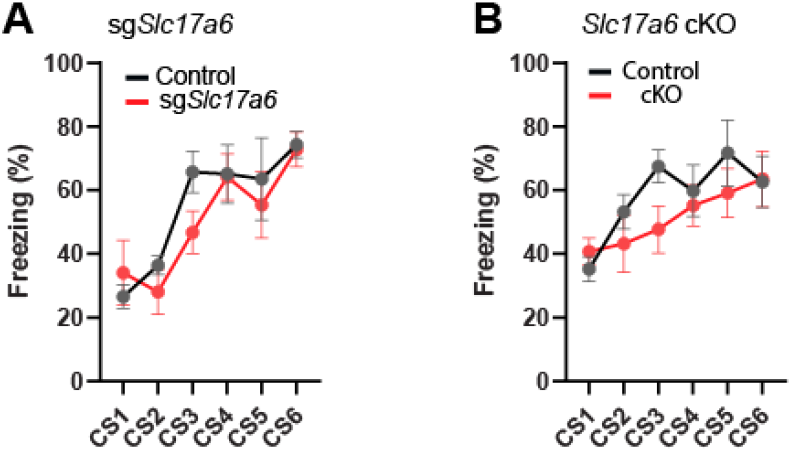
Learning curves during fear conditioning. (A) Freezing during tone (CS) in learning sessions (n = 4 for control, n = 6 for *sgSlc17a6*). (B) Freezing during tone (CS) in learning sessions (n = 7 for control, n = 6 for cKO).

